# Dynamic Changes in Endometrial Folding and Secretory Activity Across the Menstrual Cycle

**DOI:** 10.64898/2026.09.24.754255

**Authors:** Jessica Garcia De Paredes, Anat Chemerinski, Lipika Murali, Beatrice Lynch, Qingshi Zhao, Gregory Burns, Emmanuel Paul, Nataki C. Douglas, Ripla Arora

## Abstract

Embryo implantation remains a major limitation of assisted reproductive technology, with failure occurring in approximately 30% of euploid embryo transfers. Implantation requires a synchronized dialogue between the blastocyst and receptive endometrium during the window of implantation (WOI), yet minimally invasive approaches to characterize the structural and molecular features of receptivity remain limited. We analyzed paired sonohysterogram images and uterine lavage samples collected during the proliferative and mid-secretory phases from subjects with regular ovulatory cycles and proven fertility. Endometrial folds were quantified, and lavage samples were analyzed by Luminex multiplex immunoassay. Folds were present in both phases but were significantly more abundant during the mid-secretory WOI, independent of imaging view and endometrial thickness. Folding correlated strongly with circulating estradiol level during the proliferative phase but not the mid-secretory phase, and folding patterns between phases were not correlated, suggesting distinct regulatory mechanisms. Consistent with these structural patterns, uterine lavage demonstrated phase-specific differences in expression of factors associated with endometrial receptivity and implantation, with glandular epithelium, and myeloid-lineage cells emerging as major contributors. Together, these findings identify coordinated structural and secretory processes during the WOI and support further evaluation of endometrial folding and uterine lavage as complementary, minimally invasive markers of endometrial receptivity.

**Significant Statement:** This study identifies coordinated structural and secretory features of the human endometrium during the window of implantation, highlighting endometrial folding and uterine lavage as complementary, minimally invasive approaches for investigating endometrial receptivity.

## INTRODUCTION

Despite significant advances in assisted reproductive technologies embryo implantation rates remain lower than expected. The implantation rate following a euploid embryo transfer has been reported to be 60-70% (1), suggesting a failure rate of approximately 30% under optimal circumstances. Implantation requires a synchronized dialogue between the receptive endometrium and the blastocyst (2) and significant attention has been directed to uncovering factors that contribute to, or promote, endometrial receptivity (3). Recurrent implantation failure (RIF) is variably defined but most commonly described as failure to achieve a pregnancy after three embryo transfer cycles with high-quality embryos (4, 5)(6). Although the clinical definition and significance of RIF are debated, failure of chromosomally normal (euploid) embryos to implant highlights a critical gap in our understanding of successful implantation and points to an important contribution of endometrial factors in establishing pregnancy.

Endometrial receptivity has been the focus of significant attention, and endometrial receptivity testing is an active area of research. Endometrial receptivity refers to the limited time during which the endometrium, after undergoing morphologic and functional changes under the influence of estrogen and progesterone, is permissive to embryo attachment (7). It is hypothesized that there is a narrow window during which implantation is possible, termed the window of implantation (WOI). Endometrial receptivity testing is based on the premise that through imaging or molecular studies, the WOI can be characterized, abnormalities can be diagnosed, and conditions for embryo transfer can be optimized. Clinically, ultrasound imaging with assessment of endometrial thickness and echogenicity represents the most frequently used tool to assess endometrial receptivity. This approach does not have a clear predictive value with respect to embryo implantation, with some studies reporting an association between increasing thickness and higher pregnancy rates (8, 9) and others reporting none (10).

Due to limitations in existing clinical methods, ongoing research has focused on developing novel approaches to improve the assessment of endometrial receptivity. We, and others, have investigated the endometrial transcriptome to identify molecular signatures associated with the receptive state (11–13). Although earlier studies did not produce a clinically validated tool for assessing receptivity (14), recent published data from our lab (13) demonstrated that the receptivity signature is primarily driven by the glandular epithelium. This led to the development of the Glandular Epithelium Receptivity Module (GERM) score, a transcriptomic signature comprising 556 key genes that can be quantified from bulk RNA sequencing data, providing a cell type-informed approach to assessing receptivity that had not previously been explored. However, a key limitation is that transcriptomic analysis requires an endometrial biopsy, an invasive procedure that is typically not performed during a treatment cycle.

Analysis of uterine secretions collected via uterine lavage, which is a minimally invasive procedure, has also emerged as a promising strategy for identifying biomarkers of endometrial receptivity. Identified candidate biomarkers include placental growth factor (PlGF) (15), ectonucleotide pyrophosphatase/phosphodiesterase 3 (ENPP3) (16), and larger panels including desmoplakin (17). Endometrial secretions are presumed to arise from the luminal and glandular epithelium and play an important role in embryo signaling (18). Data from recurrent implantation failure suggests that glandular and luminal secretions differ in these patients with differentially abundant miRNAs (19). In many mammalian species including the mouse, leukemia inhibitory factor (LIF) and vascular endothelial growth factor A (VEFGA) are two secreted proteins that have been shown to play a crucial role in embryo implantation and angiogenesis at embryo implantation sites, respectively (20–23). LIF and VEGFA have both been detected at significantly higher levels in the WOI in uterine lavage from women and markedly reduced levels for both were detected in women with unexplained infertility (24, 25).

Analysis of endometrial structure remains an underexplored area and represents a less invasive approach to endometrial receptivity testing. Sonohysterography (SH), a minimally invasive procedure used to evaluate the presence of endometrial pathology through saline distension of the uterine cavity, represents a modality that can be used to identify structural variations within the endometrium. While most SH studies are performed in the proliferative phase, it has been noted that in the secretory phase the endometrium has an undulating pattern, with distinct structures identified as “folds” or “moguls” (26). These findings were considered pathologic until Jokubkiene et al prospectively enrolled women to undergo SH in the secretory phase, 6-10 days after a positive luteinizing hormone surge, and found that 46% (12/26) of participants demonstrated the pattern of endometrial folds previously described (27). Importantly, endometrial folds were detected independent of endometrial pathology. However, these assessments were limited to the secretory phase, and few subsequent studies have substantiated the findings. While the presence and physiological role of endometrial structural changes in humans remains unclear, our studies in rodents suggest that tissue folding is critical for embryo implantation and pregnancy success (28, 29). We have previously reported on the use of confocal imaging and 3D reconstruction from 2D optical slices to examine endometrial architecture in the mouse uterus (30). More recently, we demonstrated that in the mouse, there is a change in the orientation of endometrial folds (from longitudinal to transverse) coincident with rising progesterone levels in the WOI, and that aberrant folding patterns lead to defective embryo morphogenesis and mid-gestation pregnancy loss (28), supporting a key role for fold orientation in supporting pregnancy success. Together, these studies point to significant gaps in the existing literature on endometrial receptivity testing, and the underpinnings of implantation failure.

We undertook this study to investigate the presence of endometrial folds and secreted factors during the proliferative and mid-secretory phases of the human menstrual cycle and to explore their potential role in guiding embryo implantation. Specifically, our objectives were to characterize endometrial folds across these cycle phases using SH imaging, identify proteins present in uterine lavage fluid collected at the time of SH, and evaluate both structural and secretory features as potential markers of endometrial receptivity in the WOI. We predicted that endometrial folds would be more abundant during the mid-secretory WOI than during the proliferative phase and that uterine lavage fluid obtained during the WOI would be enriched with proteins that facilitate embryo implantation.

## MATERIALS AND METHODS

### Human subjects

Sonohysterogram and uterine lavages were performed on subjects with ovulatory, 21-to 35-day menstrual cycles. Using an Institutional Review Board (IRB)-approved protocol at Rutgers Health (Pro2022000670), self-identified Black and Hispanic women, aged 30-41 years, with regular, ovulatory menstrual cycles and at least one prior pregnancy were prospectively enrolled between August 2023 and May 2025. All study participants provided signed informed consent for the review of their medical records, performance of the sonohysterogram for evaluation of endometrial folds, and collection of blood and uterine lavage samples.

Women ages 21 – 45 who were not actively trying to conceive; had not used hormonal treatment in the 3 months prior to recruitment; and had undergone bilateral tubal ligation as a permanent sterilization procedure were eligible for the study. Women with abnormal uterine bleeding, including irregular menses, or with anatomic disorders of the reproductive tract, including hydrosalpinges, endometrial polyps, or fibroids, were excluded. The demographic characteristics of participants are presented in **(Table 1)**.

**Table 1.** Demographic characteristics of study subjects. Characteristics of study participants are summarized. BMI, body mass index; kg/m²; kilograms per square meter; G, gravidity; P, parity; E2, estradiol; P4, progesterone; pg/ml, picograms per milliliter; ng/ml, nanogram per milliliter; ET, endometrial thickness; mm, millimeter.

| Table 1. Demographic Characteristics of Study Subjects |  |  |  |  |  |  |  |  |  |  |
| --- | --- | --- | --- | --- | --- | --- | --- | --- | --- | --- |
| Subject | ID | Age | BMI (kg/m<sup>2</sup>) | Race/ ethnicity | G | P | Cycle Phase | E2 (pg/ml) | P4 (ng/ml) | ET (mm) |
| 1 | 2 | 32 | 42.07 | Black/African American | 3 | 1 | Proliferative | 24.5 | 0.1 | 5.14 |
|  | 3 |  |  |  |  |  | Mid-secretory | 145 | 9.1 | 8.97 |
| 2 | 4 | 36 | 39.57 | Hispanic or Latino | 4 | 3 | Proliferative | 100 | 0.1 | 7.1 |
|  | 5 |  |  |  |  |  | Mid-secretory | 356 | 10.3 | 10.23 |
| 3 | 6 | 35 | 41.6 | Hispanic or Latino | 3 | 3 | Proliferative | 103 | 0.1 | 11.18 |
|  | 15 |  |  |  |  |  | Mid-secretory | 168 | 10.2 | 17.56 |
| 4 | 7 | 34 | 33.8 | Hispanic or Latino | 4 | 3 | Proliferative | 87.8 | 0.1 | 7.15 |
|  | 8 |  |  |  |  |  | Mid-secretory | 246 | 19 | 11.36 |
| 5 | 9 | 32 | 39.6 | Hispanic or Latino | 5 | 4 | Proliferative | 61.5 | 0.2 | 7.32 |
|  | 10 |  |  |  |  |  | Mid-secretory | 52.1 | 6.5 | 11.34 |
| 6 | 11 | 39 | 36.4 | Black/African American | 4 | 3 | Proliferative | 45.3 | 0.1 | 7.1 |
|  | 13 |  |  |  |  |  | Mid-secretory | 194 | 16.4 | 10.91 |
| 7 | 12 | 38 | 29.1 | Hispanic or Latino | 2 | 2 | Proliferative | 138 | 0.1 | 5.5 |
|  | 21 |  |  |  |  |  | Mid-secretory | 108 | 8.3 | 7.1 |
| 8 | 14 | 39 | 41.9 | Hispanic or Latino | 3 | 3 | Proliferative | 98.6 | 0.1 | 5.66 |
|  | 25 |  |  |  |  |  | Mid-secretory | 179 | 14.8 | 15.55 |
| 9 | 16 | 35 | 27.8 | Black/African American | 3 | 3 | Proliferative | 147 | 0.1 | 7.4 |
|  | 18 |  |  |  |  |  | Mid-secretory | 397 | 26.8 | 11.9 |
| 10 | 19 | 35 | 25 | Hispanic or Latino | 4 | 4 | Proliferative | 92.7 | 0.1 | 4 |
|  | 22 |  |  |  |  |  | Mid-secretory | 162 | 8.6 | 12.2 |
| 11 | 20 | 34 | 41.2 | Black/African American | 4 | 4 | Proliferative | 121 | 0.1 | 8 |
|  | 26 |  |  |  |  |  | Mid-secretory | 145 | 10.6 | 9 |
| 12 | 23 | 41 | 32.6 | Hispanic or Latino | 7 | 5 | Proliferative | 73.1 | 0.2 | 7.3 |
|  | 24 |  |  |  |  |  | Mid-secretory | 157 | 19 | 7.9 |
| 13 | 27 | 41 | 39 | Black/African American | 6 | 6 | Proliferative | 178 | 0.1 | 6.06 |
|  | 29 |  |  |  |  |  | Mid-secretory | 192 | 17.7 | 10.49 |
| 14 | 28 | 29 | 35.7 | Hispanic or Latino | 4 | 3 | Proliferative | 56.8 | 0.1 | 4.5 |
|  | 30 |  |  |  |  |  | Mid-secretory | 64.1 | 5.3 | 12.87 |
| 15 | 31 | 34 | 39.87 | Black/African American | 4 | 2 | Proliferative | 82.9 | 0.1 | 5.1 |
|  | 32 |  |  |  |  |  | Mid-secretory | 507 | 15.5 | 10.7 |

### Uterine lavage and sonohysterography Procedures

Subjects underwent uterine lavage, followed by sonohysterogram in the proliferative phase (cycle day 6-11) and mid-secretory phase (7-10 days after detecting the urinary luteinizing hormone (LH) surge, or LH+8-10), yielding 12 paired samples for uterine lavage studies, and 15 paired samples for endometrial fold assessment.

For the uterine lavage, a one-sided speculum was placed into the vagina, and the cervix and vagina were cleansed with Betadine®. 10 mL of sterile 0.9% saline solution was infused into the uterine cavity through a fine flexible catheter aspirated, and stored in 1-1.5 mL aliquots at −80°C .

For the sonohysterogram, a transcervical balloon catheter was placed into the uterus and inflated at the level of the endocervix. The speculum was removed and a transvaginal ultrasound probe was used to visualize the uterus and adnexa. Endometrial thickness was measured. To distend the uterine cavity, 0.9% sterile saline was infused. In a sagittal orientation, images of the mid-sagittal uterus and the left and right cornua were captured. In a transverse uterine view, images of the cervix, lower uterine segment, and uterine fundus were captured. Images were saved in Viewpoint®, exported in a digital imaging and communications in medicine (DICOM) format.

### Processing uterine lavage samples

1 mL of each lavage sample was loaded onto Microsep UF spin filter columns (3 kDa molecular mass cutoff) (PALL Life Sciences, Port Washington, NY) and centrifuged at 4800g for 22 minutes to reduce the volume to 150 mL. The concentrated, eluted sample was transferred to fresh tubes. Protein concentrations were determined using the Pierce Bicinchoninic acid (BCA) Protein Assay (Thermo Fisher Scientific), according to the manufacturer’s instructions. In brief, bovine serum albumin (BSA) standards were prepared from a 2 mg/mL stock solution and serially diluted in RIPA buffer to generate a standard curve. Uterine lavage samples were diluted 1:10 in RIPA buffer. Standards and samples were assayed in duplicate, incubated at 37°C for 30 min with gentle shaking, and then cooled to room temperature. Absorbance was measured at 562 nm within 10 min using a microplate reader, and protein concentrations were calculated from the standard curve.

### Luminex multiplex assay

Concentrations of 45 human cytokines, chemokines, and growth factors in uterine lavage samples (n=12) were quantified using the Invitrogen ProcartaPlex^TM^ Luminex assay (ProcartaPlexTM^TM^ Human Cytokine/Chemokine/Growth Factor Convenience Panel 1, 45-Plex;Thermo Fisher Scientific, Waltham, MA) according to the manufacturer’s instructions.

In brief, capture bead mix (50 µL) was added to each well of a 96 well plate and washed. Universal Assay Buffer (25 µL all wells) and uterine lavage samples, standard, and background were added (25 µL per respective well), then incubated with shaking at room temperature for 2 hours to allow analyte binding. Following incubation, plates were washed and incubated sequentially with biotinylated detection antibody mix (25 µL per well) and streptavidin–phycoerythrin (50 µL per well), with washes performed between each step. After a final wash, beads were resuspended in the 1x wash buffer, and plates were analyzed. Analysis of each sample was performed in triplicate.

The lower limit of quantitation (LLOQ) was established per analyte based on the lowest qualified standard point that was greater than the limit of detection (LOD). The LOD was considered as background mean plus two standard deviations. Data were collected on the xMAP INTELLIFLEX^TM^ instrument following kit recommended setup. Analysis was performed using ProcartaPlex analysis app (Thermo Fisher Scientific, Waltham, MA).

### Quantification of endometrial folds

DICOM images were converted into .jpg or .png format. Images were filtered to include only images where the endometrium was clearly distended. Images that had the ovaries or where the endometrium was not distended were excluded from EF analysis. Images were compiled sequentially from each subject. The observer was blinded to the phase in which the US images were collected. EFs were defined as focal endometrial thickenings with shallow or deep indentations outward from the lumen into the surrounding stroma. EFs were manually annotated on each image. EFs from an individual were then compiled in an excel file. Folds per image per subject (FPIPS) were calculated. EF data was also compared against hormone levels and endometrial thickness to determine correlations.

### Serum

At each study visit, blood (5ml) was collected in nonheparinized, serum separator tubes, allowed to clot, and centrifuged at 4°C. Serum estradiol (E2) and progesterone (P4) levels were determined.

### Bulk RNA sequencing analysis

Total RNA from endometrial tissue was prepared using the KAPA Stranded RNA-Seq Kit, sequenced on an Illumina NextSeq 500 (GSE289073), and mapped to the GRCh38 genome using STAR (v2.7.9a). Reads overlapping Ensembl (v110) annotations were counted. Differential expression was evaluated using the edgeR-robust method, filtering out low-count genes via filterByExpr. Genes were considered significantly differentially expressed if the FDR-corrected p-values were less than 0.05.

### Single-cell RNA sequencing (scRNA-seq) analysis

Single-cell libraries were generated using the 10X Chromium Single Cell 3′ GEM kit (v2) and sequenced on an Illumina NextSeq 500 (GSE290822). Reads were processed with Cell Ranger (v3.1.0) and analyzed using Seurat (v5.1.0) in R. Cells expressing fewer than 200 genes or with >5% mitochondrial content were excluded. Following dataset integration, data normalization (LogNormalize), and dimensionality reduction using the top 20 principal components, cell clustering was performed. The Seurat DotPlot function was utilized to visualize the gene expression of the Luminex cytokines within the mid-secretory phase samples. The Seurat function AddModuleScore was used to calculate the combined gene signature score for CCL2, CCL3, CXCL10, IL6, LIF, VEGF-A. The full details of the sequencing technique and analysis are described in our previous publication (13).

### Statistical analysis

For all analyses, a Shapiro-Wilk test was used to determine normality of the dataset. Normally distributed data were expressed as a mean with standard deviation (SD) and non-normally distributed data were expressed as a median with interquartile range [IQR]. A paired t test or Wilcoxon matched pairs signed rank test was used to compare continuous variables. The Spearman correlation coefficient was calculated to determine the association between FPIPS and menstrual cycle phase, endometrial thickness, serum E2 or serum P4 level. *p*<0.05 was considered statistically significant for all analyses. Statistical analyses were performed with Prism v11.0.2 (GraphPad Software).

## RESULTS

### Demographic data and experimental study design

The mammalian endometrium undergoes cyclic remodeling across the menstrual/estrus cycle in preparation for embryo implantation. To assess gross morphological changes in the human endometrium and changes in the uterine secretome across the menstrual cycle, we enrolled participants with regular ovulatory menstrual cycles and proven fertility. In our cohort, 40% of participants self-identified as Black/African American (6/15) and 60% (9/15) as Hispanic. Participants in the study had a mean +/- SD age of 35.3 +/- 3.7 years and a mean +/- SD BMI of 35.7 +/- 5.6 kg/m2. The mean +/- SD gravidity and parity were 4.1 +/-1.2 prior pregnancies and 3.1 +/- 1.0 live births, respectively. Characteristics of the study participants are summarized in **Table 1**.

After enrollment, study participants underwent uterine lavage and SH during the proliferative phase (cycle days 6-11) and the mid-secretory WOI, which was defined as 8-10 days after detection of a LH surge in urine, yielding a total of 15 paired samples for endometrial fold assessment and 12 paired samples for uterine lavage studies. Serum E2 and P4 levels and endometrial thickness were determined at each study visit. **(Table 1** and **Figure 1, Panel A)** The median [IQR] serum E2 level was significantly lower during the proliferative phase than during the mid-secretory phase (90.3 [58.0, 116.5] vs. 165.0 [117.3, 233.0] pg/mL, p = 0.0003). As expected, the median [IQR] serum P4 level was also significantly lower during the proliferative phase than during the post-ovulatory, mid-secretory WOI (0.1 [0.1, 0.1] vs 10.6 [8.6, 17.7] ng/mL, p = 0.0001). Likewise, the mean +/- SD endometrial thickness was significantly lower during the proliferative phase than in the mid-secretory phase [6.5 +/- 1.7 vs 11.2 +/- 2.7 mm, p < 0.0001]. Together, these data confirm the expected hormonal and sonographic differences between the proliferative and mid-secretory phases of the menstrual cycle.

**Figure 1.**
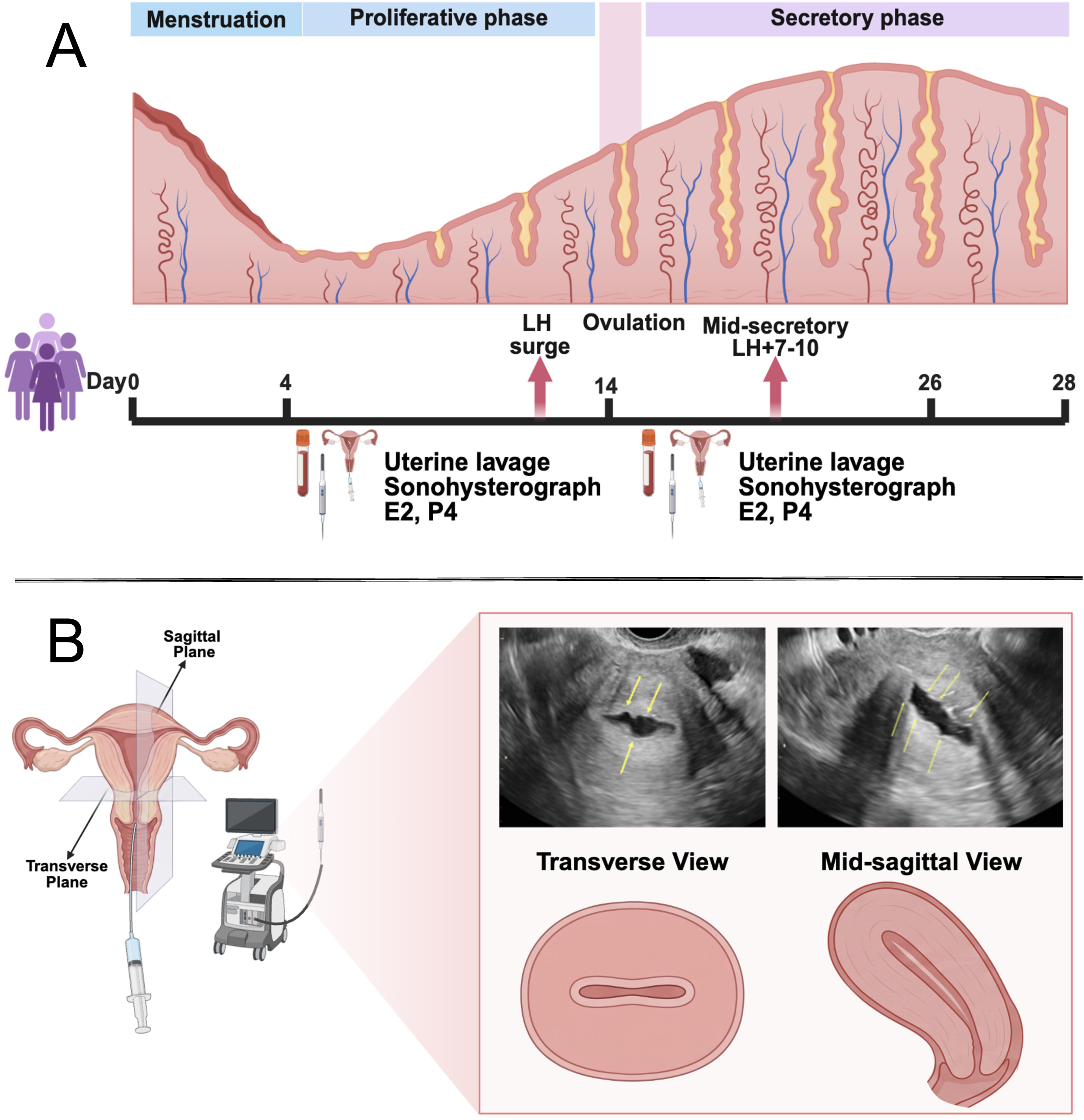
Flowchart of the study design and endometrial folds detection. (A) Schematic representation of enrolled subjects and sample collection. (B) Schematic illustration of the sonohysterography procedure and uterine lavage sample collection. Yellow arrowheads indicate endometrial folds visualized on transverse (left) and mid-sagittal (right) sonohysterographic ultrasound images. LH, luteinizing hormone; E2, estradiol; P4, progesterone.

### Endometrial folds are increased during the window of implantation

Utilizing SH, an evaluation of the endometrial gross morphology was performed to characterize EFs in the proliferative phase and mid-secretory WOI. SH images were evaluated for the presence and number of outward EFs. EFs were observed in both sagittal and transverse views; both shallow and deep indentations were observed in subjects from proliferative and mid-secretory phases of the menstrual cycle. **(Figure 1, Panel B**). Quantification of EFs showed an average of 1.7 +/- 0.7 folds per image per subject (FPIPS) in the proliferative phase and 3.2 +/- 0.9 FPIPS in the mid-secretory phase, resulting in a statistically significant, menstrual phase-specific difference **(Figure 2, Panel A).**

**Figure 2.**
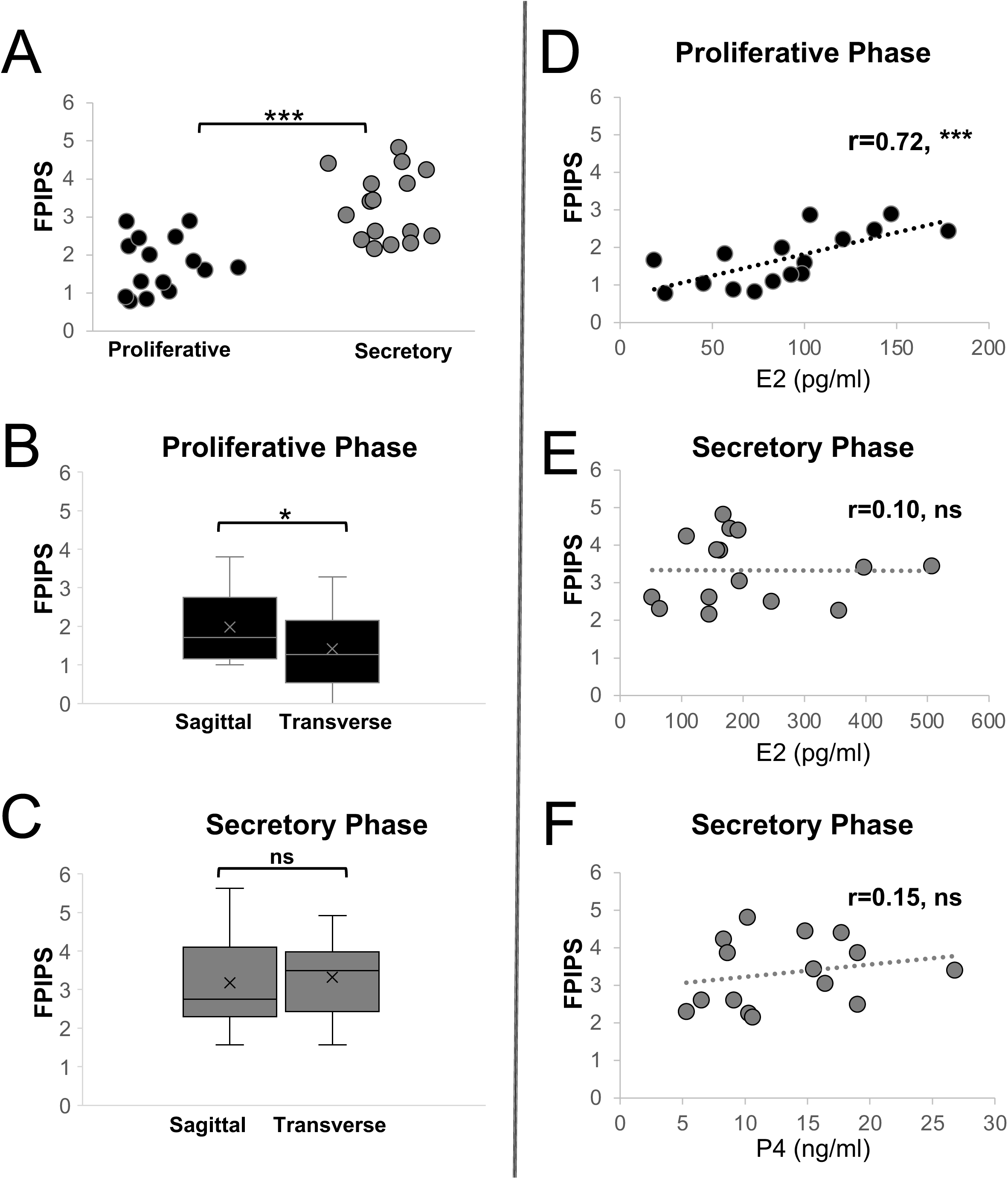
Endometrial folds are increased during the window of implantation. (A) Quantification of endometrial FPIPS during the proliferative and secretory phases. Statistical analysis was performed using paired two-tailed t-test. (B-C) Quantification of mean endometrial FPIPS folds in sagittal versus transverse US view during the proliferative and secretory phases. Data are presented as median (IQR). Statistical analysis was performed using the Wilcoxon signed-rank test. (D-F) Linear regression analyses demonstrating the relationship between hormone levels and the number of endometrial FPIPS across menstrual cycle phases. Statistical analysis was performed using Spearman correlation coefficient and presented as the coefficient of determination (r). \**p* < 0.05, \*\*\**p* < 0.001. FPIPS, folds per image per subject; US, ultrasound; E2, estradiol; P4, progesterone; pg/ml, picograms per milliliter, ng/ml, nanogram per milliliter. NS, not statistically significant.

The uterine lining can be folded along the rostral-caudal axis (captured in a transverse view of the ultrasound) or the dorsal-ventral or medial-lateral axis (captured in the sagittal view of the ultrasound) **(Figure 1, Panel B)**. To determine axes of endometrial folding we evaluated if the folds observed in each of the phases were predominantly found in sagittal or transverse US views. For the proliferative phase when evaluating FPIPS by plane there was a significant increase in folds in the sagittal US views when compared to transverse views (p=0.03). For the secretory phase there was no significant difference in FPIPS observed in transverse or sagittal US views **(Figure 2, Panel B-C)**.

Given that endometrial thickness is a well-documented physiological change that varies with phase of the menstrual cycle, the association of EFs with endometrial thickness was determined **(Supplemental Figure 1, Panel A)**. A positive correlation was observed with the number of FPIPS and endometrial thickness in the proliferative phase (r=0.33, p=0.21) however the association was not statistically significant. There is no correlation between the FPIPS and the endometrial thickness in the mid-secretory phase (r=0.06, p=0.8) **(Supplemental Figure 1, Panel A),** suggesting that an increased amount of endometrial tissue is not responsible for the increased number of EFs observed in the mid-secretory phase.

### Presence and number of endometrial folds are hormonally regulated

To assess the association of EFs with circulating levels of ovarian-derived hormones, the extent of folding was correlated with menstrual cycle phase-specific serum E2 levels and mid-secretory phase P4 levels. P4 levels were not analyzed during the proliferative phase because, in the absence of a post-ovulatory corpus luteum, serum P4 concentrations were low (0.1 - 0.2 ng/mL), with a range that is not biologically meaningful for correlation analyses. In the proliferative phase, the FPIPS showed a strong positive correlation with serum E2 levels (r=0.72, p=0.002), suggesting that endometrial folding during the proliferative phase is driven by the levels of circulating estrogen **(Figure 2, Panel D).** In contrast, during the mid-secretory phase, endometrial folding was not correlated with serum levels of E2 (r=0.10, p=0.72) or P4 (r=0.15, p=0.59) **(Figure 2, panel E-F)**. Taken together, these findings and the greater number of folds in the mid-secretory as compared to the proliferative phase suggest that high progesterone exposure, rather than circulating progesterone level, is the principal determinant of endometrial folding during the mid-secretory phase. Notably, a subject that showed a higher number of folds in the proliferative phase did not necessarily/correlatively show a higher number of folds in the mid-secretory WOI, suggesting independent mechanisms of folding in each phase of the cycle **(Supplemental Figure 1, Panel B)**

### Phase-specific changes in protein secretion and enrichment of mRNAs encoding the uterine secretome during the window of implantation

In addition to structural changes, the mid-secretory WOI is characterized by global changes in gene expression, protein regulation, and protein secretion. To evaluate these variations across the proliferative and mid-secretory phases, uterine secretome analysis was performed on uterine lavage samples to identify detectable secreted proteins and assess the presence of factors that could support embryo implantation.

First, total protein concentration in the proliferative and mid-secretory phase was determined. In the proliferative phase, the total protein concentration ranged from 203.8 to 4034 ng/mL and in the mid-secretory phase the total protein concentration ranged from 203.8 to 1938 ng/mL. The median total protein concentration was significantly higher in the proliferative phase compared to the mid-secretory WOI (1024 [572.8, 2667] vs 565.7 [341.9, 1302] ng/mL, p=0.0098,) **(Supplemental Table 1)** However, for 3 of 12 subjects, the total protein concentration was higher in the mid-secretory WOI as compared to the proliferative phase.

Lavage fluid was then used to determine the concentration of 45 factors, including cytokines, chemokines, and growth factors, using the Invitrogen ProcartaPlex Luminex assay. The manufacturer-preconfigured panel was selected based on prior published studies analyzing uterine lavage samples that reported reliable detection of the included analytes (29, 31–33) **(Table 2)**

**Table 2.** Proteins secreted and mRNAs expressed in the endometrium. Summary of proteins detected in uterine lavage samples and their corresponding mRNA transcript abundance in endometrial biopsies as measured by bulk RNA sequencing across menstrual cycle phases. Uterine lavage values are presented as median (25th–75th percentiles). Bulk RNA sequencing results are reported as log₂ fold change (log₂FC) and false discovery rate (FDR). CCL, chemokine ligand; CXCL, chemokine (C-X-C motif) ligand; IL, interleukin; LIF, leukemia inhibitory factor; PLGF, placental growth factor; VEGF, vascular endothelial growth factor; FGF2, fibroblast growth factor 2;HGF, hepatocyte growth factor; PDGF-BB, platelet-derived growth factor BB; SDF-1 alpha, stromal cell-derived factor 1-alpha.

| Table 2. Proteins secreted and mRNAs expressed in the endometrium |  |  |  |  |  |  |  |
| --- | --- | --- | --- | --- | --- | --- | --- |
| Protein | Gene Symbol | Concentration of Mediator (median;pg/ml) Luminex |  |  |  | Bulk RNA |  |
|  |  | Proliferative | Secretory |  |  | Mid-secretory vs Proliferative |  |
|  |  |  |  | FC | p-value | FC | FDR p-value |
| MCP-1 | CCL2 | 160.0 | 321.2 | 2.00 | 0.004 | 1.46 | 4.52 |
| MIP-1 alpha | CCL3 | 3.505 | 17.47 | 4.98 | 0.001 | 2.06 | 1.93 |
| Eotaxin | CCL11 | 0.7550 | 4.110 | 5.44 | 0.0005 | N/A | N/A |
| IP-10 | CXCL10 | 13.08 | 104.2 | 7.96 | 0.0005 | 0.90 | 1.34 |
| IL-6 | IL-6 | 24.79 | 169.7 | 6.84 | 0.25 | 2.13 | 6.78 |
| LIF | LIF | 6.230 | 34.58 | 5.55 | 0.01 | 3.51 | 3.55 |
| PLGF-1 | PGF | 3.880 | 11.33 | 2.92 | 0.002 | -0.97 | 2.39 |
| VEGF-A | VEGFA | 30.86 | 470.1 | 15.23 | 0.001 | 0.70 | 1.52 |
| VEGF-D | VEGFD | 0.26 | 0.86 | 3.30 | 0.07 | N/A | N/A |
| MIP-1 beta | CCL4 | 23.59 | 57.50 | 2.43 | 0.06 | 1.33 | 2.58 |
| RANTES | CCL5 | 5.040 | 4.665 | 0.92 | 0.96 | -0.12 | 8.04 |
| GRO alpha | CXCL1 | 133.4 | 311.4 | 2.33 | 0.003 | 1.86 | 1.00 |
| FGF-2 | FGF2 | 265.5 | 45.65 | 0.17 | 0.64 | -0.05 | 8.69 |
| HGF | HGF | 195.5 | 171.8 | 0.87 | 0.62 | -0.56 | 1.65 |
| IL-8 | CXCL8 | 105.9 | 294.8 | 2.78 | 0.002 | 1.32 | 1.03 |
| IL-18 | IL-18 | 5.680 | 9.445 | 1.66 | 0.25 | 0.33 | 3.55 |
| PDGF-BB | PDGFB | 5.850 | 10.35 | 1.76 | 0.14 | -0.06 | 8.47 |
| SDF-1 alpha | CXCL12 | 67.49 | 87.04 | 1.28 | 0.20 | -0.78 | 9.75 |

Uterine lavage secretome analysis revealed that of the 45 analytes assessed, 18 proteins were detected in more than 95% of uterine lavage samples in both the proliferative and the mid-secretory phases. These include CCL2, CCL3, CCL4, CCL5, CCL11, CXCL1, CXCL10, FGF2, HGF, IL-6, IL-8, IL-18, LIF, PDGF-BB, PLGF-1, VEGFA, VEGFD, and SDF-1alpha **(Figure 3, Panel A** and **Supplemental Figure 2)**. Concentrations of detected factors from paired lavage samples were compared between menstrual cycle phases using the Wilcoxon signed-rank test and phase-specific enrichment of factors was observed. Concentrations of CCL2, CCL3, CCL11, CXCL10, IL-6, LIF, PLGF-1, VEGFA, and VEGFD were significantly increased during the mid-secretory phase compared to the proliferative phase. (**Figure 3, Panel A)** None of the factors were significantly increased in the proliferative phase as compared to the mid-secretory phase (**Supplemental Figure 2**).

**Figure 3.**
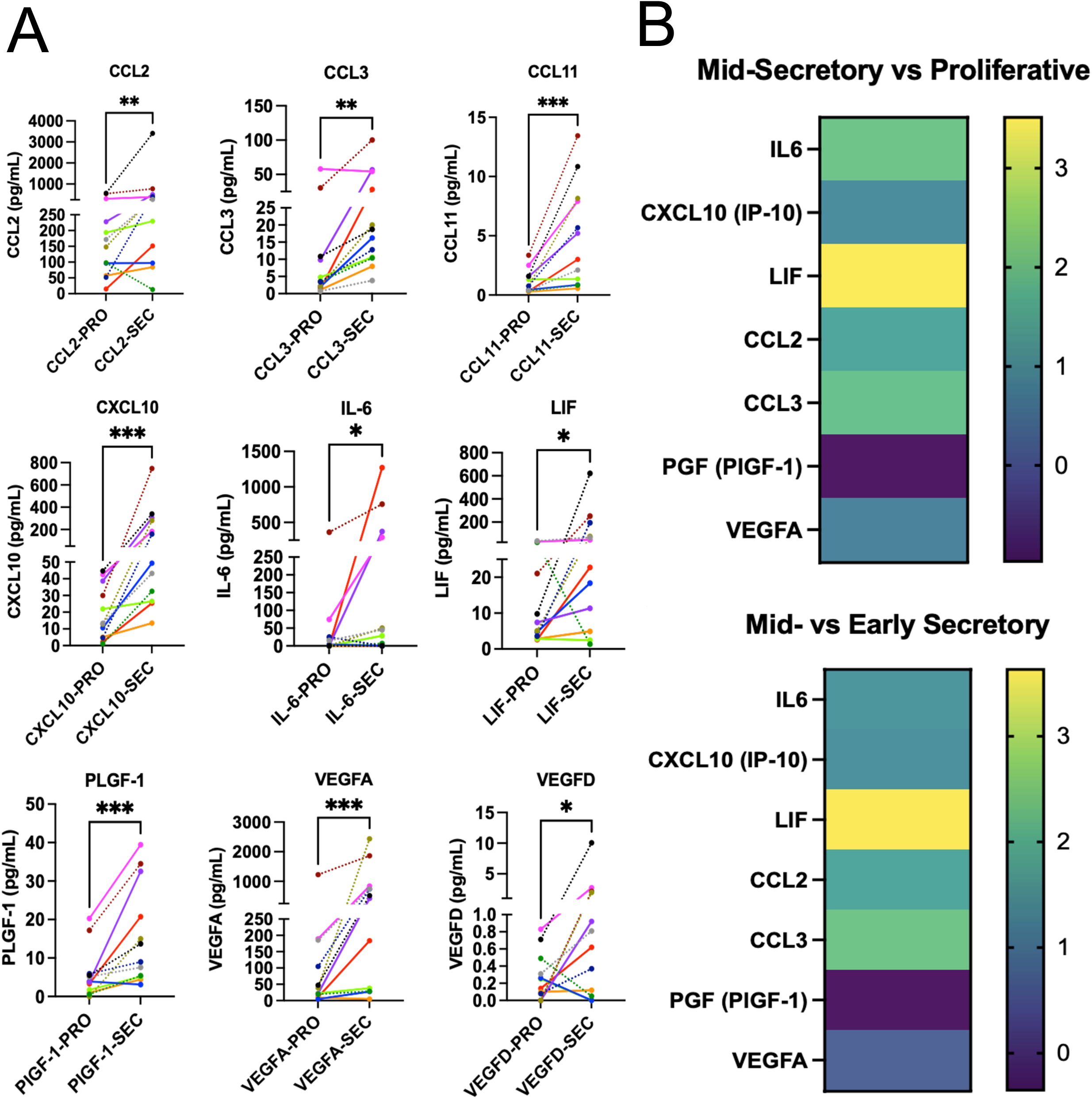
Phase-specific changes in protein secretion and enrichment of mRNAs encoding the uterine secretome during the window of implantation. (A) Comparison of proteins significantly increased in uterine lavage samples during the mid-secretory WOI phase relative to the proliferative phase. Data are presented as median (IQR). Statistical analysis was performed using the Wilcoxon signed-rank test, with \**P* < 0.05, \*\**P* < 0.01, \*\*\**P* < 0.001. (B) Heat map comparing the mRNAs for the proteins enriched in lavage during the WOI from bulk RNA sequencing of endometrial biopsies. Mid-secretory vs proliferative phase (top) and mid-secretory vs early secretory phase (bottom). Colors represent the relative expression of a factor; yellow indicates an increased expression and purple a decreased expression. WOI, window of implantation. CCL, chemokine ligand; CXCL, chemokine (C-X-C motif) ligand; IL, interleukin; LIF, leukemia inhibitory factor; PLGF, placental growth factor; VEGF, vascular endothelial growth factor.

### Bulk RNA and single cell RNA sequencing analysis

To ascertain if secreted factors enriched in the uterine lavage correlate with global changes in the endometrial transcriptome across the menstrual cycle, we evaluated bulk mRNA-seq data from proliferative and secretory phase endometrial tissue (13). We then identified the cell types expressing the 45 factors included in the ProcartaPlex Luminex panel using single cell RNA-sequencing (scRNA-Seq) data (13). Uterine lavage protein expression was compared to mRNA expression in independent cohorts.

Out of the 9 factors that were significantly enriched in uterine lavage during the mid-secretory phase, 6 factors (CCL2, CCL3, CXCL10, IL6, LIF, VEGFA) were enriched at the mRNA level in endometrial tissue samples collected during the mid-secretory WOI **(Figure 3, Panel B).** To our surprise, PlGF-1 was not differentially expressed at the mRNA level between the proliferative and the secretory phase, despite high levels of protein detected in the lavage. This could indicate a higher rate of mRNA translation in the secretory phase compared to the proliferative phase. Additionally, 2 of the 9 factors (CCL11, VEGFD) enriched in the mid-secretory phase lavage were not found in the bulk RNA seq data. These 2 factors represent possible secretion from other sources (eg. cervical) **(Figure 3, Panel B)**

Next, we analyzed scRNA-Seq data from the mid-secretory phase of the cycle to determine which cell-types express the mRNA encoding the secretory proteins found to be enriched in the mid-secretory phase uterine lavage **(Figure 4, Panel A)**. Similar to the bulk RNA-seq data we only identified expression of 6 of the factors. As expected, the endometrial glandular epithelium was identified as the source for 3 factors, including IL6, LIF and VEGFA. Surprisingly, the myeloid lineage was found to be a cell type with high mRNA expression of 5 out of the enriched 6 factors: IL6, CXCL10, CCL2, CCL3 and VEGFA (**Figure 4, Panel B** and **Supplemental Figure 3).** Other factors (9) that were detected but not increased during the mid-secretory phase were also expressed by the glandular epithelium and the myeloid lineage amongst other cell types. Taken together, the comparison of uterine lavage protein expression with endometrial mRNA expression across independent subjects demonstrated consistent expression across individuals.

**Figure 4.**
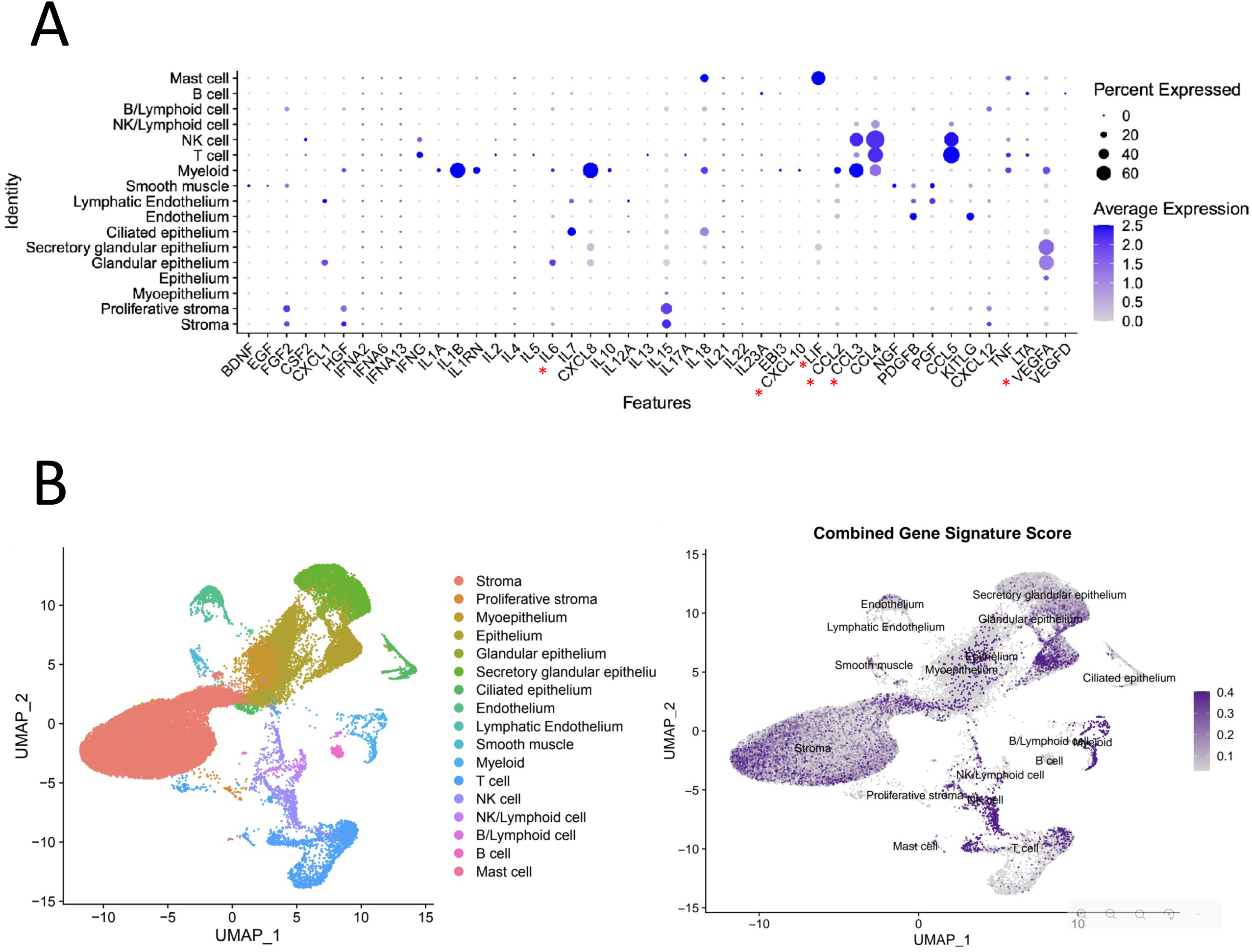
Enriched factors during the window of implantation are predominantly expressed by the myeloid lineage and the glandular epithelium. (A) Single-cell RNA sequencing analysis of endometrial samples from the mid-secretory phase. Dot plots illustrate abundance of mRNA transcripts for luminex panel proteins expressed across endometrial cell types during the mid-secretory phase. Average gene expression and the proportion of cells expressing each gene within a cell cluster are indicated by color intensity and dot size, respectively. (B) UMAP visualization of the cell-types (left) and combined gene signature score for 6 factors, CCL2, CCL3, CCL11, CXCL10, IL6 and VEGFA, enriched during the mid-secretory WOI. Combined score is represented by color intensity. WOI, window of implantation.

## DISCUSSION

Despite advances in assisted reproductive technology, implantation failure is a significant clinical challenge, occurring in approximately 30% of cycles (1). To identify structural changes and secreted factors associated with embryo implantation and successful pregnancy outcomes, we characterized sonohysterography images and uterine lavage samples during the proliferative and mid-secretory phases of the menstrual cycle from subjects with regular ovulatory cycles and proven fertility. The presence of endometrial folds in both phases was confirmed on sagittal and transverse ultrasound views. Consistent with our hypothesis, the average number of folds was significantly greater during the progesterone-dominant mid-secretory phase, independent of view and endometrial thickness. Folding was strongly associated with circulating estradiol level during the proliferative phase but not with estradiol or progesterone during the mid-secretory phase. Because our analysis was performed on paired samples, the lack of correlation between endometrial folding in the proliferative and secretory phases strongly suggests that folding is driven by independent mechanisms across the menstrual cycle. Similarly, analysis of uterine lavage samples demonstrated phase-specific differences, with enrichment of novel factors as well as factors known to be associated with endometrial receptivity and implantation, with glandular epithelium and myeloid-lineage cells emerging as major contributors to their expression. Together, these findings demonstrate coordinated structural and molecular changes during the mid-secretory window of implantation that may contribute to endometrial receptivity and successful embryo implantation.

Currently utilized methods to clinically assess endometrial receptivity during a conception cycle are non-invasive but have limited predictive value, highlighting both an incomplete understanding of receptivity and a lack of robust technologies for its assessment. For individuals undergoing fertility treatment, 2D ultrasound, in combination with serum estradiol and progesterone measurements, remains the most widely used and clinically validated approach for evaluating the endometrium during a conception cycle. Ultrasound assessment typically focuses on endometrial thickness and pattern immediately prior to ovulation or initiation of exogenous progesterone. A trilaminar endometrium measuring ≥7 mm has been associated with higher clinical pregnancy rates in some, but not all, studies (9, 34). Additional parameters that can be assessed using 2D or 3D ultrasound include endometrial blood flow by Doppler, endometrial volume, and uterine/endometrial peristalsis (35–37). However, these measures have not been validated as predictors of endometrial receptivity are therefore not routinely employed in clinical practice.

Importantly, none of the existing ultrasound-based assessments evaluates endometrial architecture, which is known to contribute to receptivity in the mouse (28). By analyzing 3D structure of the mouse uterus during early pregnancy we have established that dynamic remodeling of uterine epithelium is critical for successful embryo implantation. During the pre-implantation stage (gestational day (GD) 3.5), uterine longitudinal folds that run along the oviductal-cervical axis, transition to transverse folds that run along the mesometrial-anti-mesometrial axis. This transition is key as a failure to remove all the longitudinal folds results in embryos being trapped in folds preventing a proper implantation chamber from forming. This embryo trapping results in failure of embryo rotation, mis-alignment of the embryo’s inner cell mass-trophoblast axis to the uterine mesometrial-anti-mesometrial axis (28, 38) and mid-gestation embryo resorption and pregnancy loss (28, 31, 39). Scanning electron microscopy analysis during early pregnancy in rats revealed dynamic changes in epithelial architecture mirroring those observed in mice (33). In mares, in the estrogen rich phase, an endometrial cartwheel folding pattern indicative of edema is observed. Hyperedema is characterized by loss of the cartwheel pattern and is indicative of endometritis, cervical incompetence, aberrant vascular response to estrogens, a lymphatic pathology or alterations in myoelectric activity representing a pathological state of the endometrium (40). As progesterone levels rise the cartwheel pattern is lost during normal pregnancy. Failing to lose the cartwheel pattern or regaining the cartwheel pattern is a sign of low progesterone or endometrial infection that often results in pregnancy loss (32). Thus, endometrial remodeling is a critical indicator of early pregnancy progression.

Analysis of endometrial architecture to identify structural features associated with receptivity in the human endometrium remains relatively understudied. A recent report described the use of hysteroscopic image-recognition technology to quantify endometrial gland density and size, as well as the number and morphology of pinopodes during the window of implantation (41). Higher endometrial gland density and a greater number of pinopodes, particularly those exhibiting mature morphology, were associated with successful embryo implantation. To our knowledge, endometrial folding as an architectural feature associated with human endometrial receptivity has been examined in only one prior study which identified folds in the secretory phase without a correlation to the endometrial thickness (27).

The human proliferative phase uterus is characterized by high levels of epithelial cell proliferation, increasing endometrial thickness, and a relatively quiescent myometrium (42, 43). In tubular organs where an epithelial tube is surrounded by a smooth muscle tube such as the uterus, folding is a function of the epithelial length compared to the smooth muscle length. Further, the amount of compression applied by the smooth muscle tube can alter patterns of epithelial folding in conjunction with or independent of the effects of proliferation. In the m9kouse oviduct, the ratio of circular epithelial length to circular smooth muscle ratio is about four to one and this mechanical constraint drives epithelial fold formation (44). Thus, the increase in folds in the human proliferative phase endometrium suggests an increase in epithelial cell number resulting in larger amounts of epithelium packaged in a constrained muscular tube by the process of buckling morphogenesis (45).

Hormonally, the secretory phase is considered “progesterone dominant”, with peak levels of progesterone coinciding with the window of implantation (46). The number of folds in the mid-secretory phase did not correlate with absolute levels of estradiol or progesterone. Progesterone levels do not alter epithelial proliferation (47). However, recent work from our lab and others suggests that progesterone-dominant stages are characterized by increased myometrial contractility during the estrous cycle and in early pregnancy in the mouse (48). Further, the progesterone-dominant mid-secretory stage also displays transcriptional changes in the stromal and extracellular matrix (ECM) composition that can alter the compressive forces on the epithelium (13). Thus, epithelial folding in the human mid-secretory phase is likely driven by excess compression of the myometrium or stromal ECM changes. Previous literature suggests regulation of epithelial folding by stromal signaling in the intestine and by muscle contractions and organization in both the intestine (49–51) and the oviduct (44). How compression due to contractions of ECM regulates uterine epithelial folding will be a subject of future investigations.

Endometrial biopsy, although invasive and therefore typically not performed in conception cycles, is the most commonly used approach for sampling the endometrium to investigate the cellular and molecular changes associated with a receptive endometrial state. We and others have performed transcriptomic and proteomic analyses of endometrial tissue to identify molecular signatures associated with receptivity (11, 52–54). Additionally, studies have characterized the composition of the endometrial microbiota and its potential contribution to endometrial receptivity (55, 56). Together, these studies have led to the development of molecular endometrial receptivity tests that have been introduced into clinical practice. The invasive nature of endometrial biopsy limits its use during conception cycles, and, more importantly, there is insufficient evidence that use of these tests improves pregnancy outcomes. The most recent clinical practice guidelines do not recommend routine use of these tests to improve the likelihood of embryo implantation, including in individuals with recurrent implantation failure (57, 58).

Analysis of uterine secretions collected through minimally invasive uterine lavage allows for identification of proteins produced and secreted by the endometrium (59) without requiring an endometrial biopsy and represents a promising approach for identifying minimally invasive biomarkers of endometrial receptivity. Among the 45 analytes examined in this study, 18 proteins were detectable in both the proliferative and mid-secretory phases; the concentrations of 9 of these were significantly increased during the mid-secretory phase compared to the proliferative phase, suggesting a potential role in endometrial preparation for implantation. In a previous study by Hannan et al. (24), a similar analysis of uterine lavage was performed and 30 analytes were detected in both proliferative and secretory phase uterine lavages. 11 analytes were common between the previous study and our study: CCL2, CCL3, CCL4, CCL5, CCL11, CXCL1, CXCL10, FGF2, VEGFA, IL-6, PDGF-BB. However, of these 11 analytes, Hannan et al showed that VEGFA levels were significantly different between the proliferative and mid-secretory phase. In addition to VEGFA, our study demonstrates significant differences between the cycle phases for CCL2, CCL3, CCL11, CXCL10 and IL6. These differences could be because the previous study used a smaller sample size of 4 whereas our study used paired samples from the same subject and had a larger sample size of 12 samples in both stages. Many of these proteins have been characterized in the human endometrium and their impact studied in genetic mouse models.

Among the factors identified in our study, several belong to the IL-6 cytokine family. Consistent with the findings in our study, increased endometrial gene expression of IL-6 has also been demonstrated in the window of implantation (7), and decreased expression was observed in subjects with recurrent pregnancy loss (60) and recurrent implantation failure (61). LIF is a member of the IL-6 family and a critical regulator of endometrial receptivity and embryo implantation. We and others have found that LIF expression peaks during the mid-and late secretory phases of the menstrual cycle (60), and can be found in both the glandular epithelium and stroma (62). LIF expression in the glandular epithelium is significantly reduced in women with recurrent implantation failure (62) or unexplained infertility (63). One study that analyzed uterine flushings collected during the window of implantation found reduced levels of LIF and glycoprotein 130 (gp130), the signal-transducing component of the heterodimeric LIF receptor (LIFR), in women with unexplained infertility (64). The functional importance of LIF suggested by these clinical findings is further substantiated by mouse models. LIF is encoded by gland-specific genes, and conditional loss of *LIF* in the mouse luminal epithelium leads to implantation failure. Blastocysts recovered from the mutant mice implant normally when transferred to wild-type recipients, demonstrating an endometrial requirement for LIF (65). In contrast, implantation proceeds and fertility is preserved in mice with stromal cell-specific loss of *LIF* or *LIFR* (66). These studies support the important role of the IL-6 family in blastocyst attachment and implantation (67) and highlight important gaps in knowledge.

Beyond cytokine-mediated signaling, our findings also implicate VEGFA, a pro-angiogenic factor, in mid-secretory endometrial function. Expression of VEGFA has also been characterized in the human endometrium and its role elucidated with genetic mouse models. VEGFA is strongly expressed in the human endometrium throughout the menstrual cycle in both epithelial and stromal cell compartments, with reduced expression in the mid-secretory phase in women with recurrent pregnancy loss (68). Treatment of primary endometrial epithelial cells with human chorionic gonadotropin leads to increased production of VEGFA, highlighting the additional role of VEGFA in mediating implantation and placentation (69). Conditional loss of VEGFA using a progesterone-receptor Cre driver leads to abnormal implantation site angiogenesis, while treatment of pregnant mice with VEGF Trap (a VEGF neutralizer) leads to embryo resorption (70). VEGF signaling via its receptors, VEGFR1, VEGFR2 and VEGFR3 is also a critical mediator of post-implantation vascular remodeling in mice (71).

The concordance between detecting proteins in uterine lavage and transcriptomic upregulation during the mid-secretory phase in independent cohorts of subjects, suggests that these factors are actively produced and secreted during the window of implantation, supporting their potential biological relevance to implantation. The overlap between our findings and prior studies demonstrating reduced expression of these factors in women with recurrent implantation failure and recurrent pregnancy loss further supports the likelihood of an important role in endometrial receptivity. Continued investigation of these and other proteins in the mid-secretory uterine lavage may improve our understanding of the molecular mechanisms underlying successful implantation and identify potential targets for intervention in women with impaired fertility.

This study has several notable strengths. The cohort included self-identified Black and Hispanic women with proven fertility and no known uterine pathology, a population of “healthy subjects” that remains underrepresented in the scientific literature. Subjects served as their own controls, enabling paired comparisons between the proliferative and secretory phases while minimizing interindividual variability. Menstrual cycle phase was determined by prospective cycle tracking and use of urinary ovulation predictor kits and then confirmed by serum estradiol and progesterone concentrations. Sequential sonohysterography imaging acquisition, serum hormone measurements, and uterine lavage collection during both phases enabled direct correlation of hormonal changes with endometrial thickness, endometrial folding, and protein profiles. Lastly, uterine lavage factors were measured in triplicate and normalized to the total protein concentration, enhancing the rigor and reproducibility of the analyses.

These findings should be interpreted in the context of several considerations. The relatively small sample size may have limited power to detect subtle associations, while manual endometrial fold annotation introduces potential observer variability and highlights the need for automated methods of image analysis. Secretome analysis was restricted to a predefined panel of 45 cytokines, chemokines, and growth factors, potentially overlooking other proteins involved in endometrial receptivity and embryo implantation. Finally, some discrepancies between uterine lavage proteins and endometrial mRNA expression may reflect the complex composition of lavage specimens, including contributions from non-endometrial sources such as cervical mucus, as well as inherent differences between transcriptomic and proteomic analyses.

Our study demonstrates distinct changes in uterine architecture and secretory capacity as the endometrium prepares for implantation. A better understanding of these changes could provide missing rationale for early reproductive failures, including unexplained infertility, implantation failure and recurrent pregnancy loss. In the long-term, developing automated machine learning-based methods to quantify endometrial folding from ultrasound images and designing a custom panel to detect mid-secretory phase enriched cytokines could aid in optimizing embryo transfer protocols in the clinic.

## ACKNOWLEDGEMENTS

We thank Jim M. Giron (Thermofisher) for help with luminex data analysis, Dr. Michael Saad and Dr. Kristy Blackledge for subject recruitment, and Tracy Wu for assistance with uterine lavage sample processing.

## AUTHOR CONTRIBUTIONS

RA and ND conceptualized the study and designed the experiments. JGP, LM, BL, QZ, and AC performed the experiments. ENP, GWB, AC, NCD and RA validated the data and performed the analyses. JGP, AC, ND and RA prepared the figures and wrote and edited the manuscript. All authors reviewed and accepted the final version of the manuscript.

## DATA AVAILABILITY

All data supporting the findings of this study are available within the article or its supplementary data.

---

**Supplemental Figure 1.**
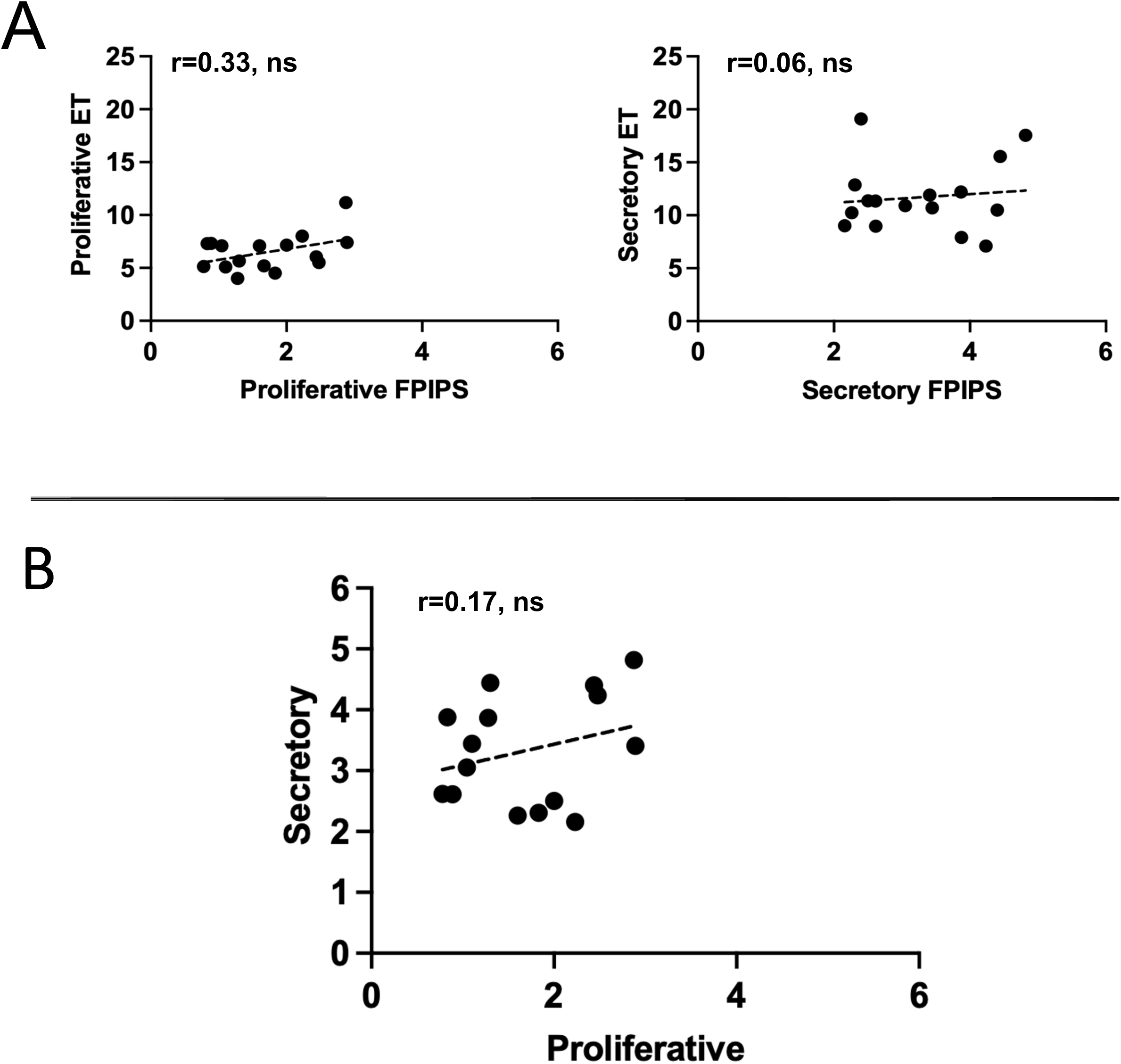
Endometrial folding is independently regulated in menstrual cycle phases. (A) Linear regression analyses demonstrates no relationship between the number of FPIPS and the endometrial thickness in the proliferative and mid-secretory phase. (B) Linear regression analyses demonstrating no relationship between FPIPS during the proliferative phase vs mid-secretory WOI phase. Statistical analysis was performed using Spearman correlation coefficient. Correlations are presented as the coefficient of determination (r). NS, not statistically significant. WOI, window of implantation; ET, endometrial thickness; FPIPS, folds per image per subject.

**Supplemental Figure 2.**
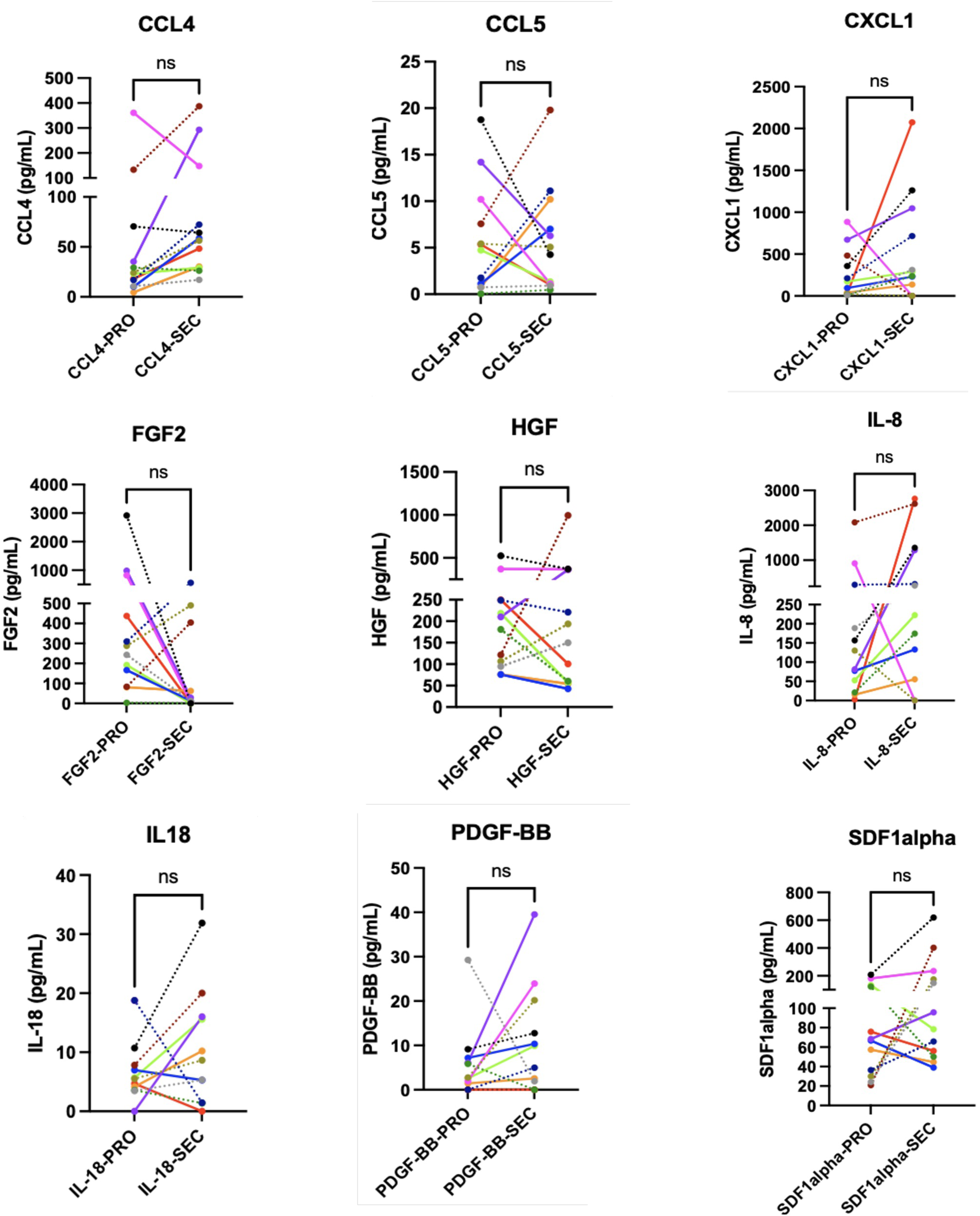
Proteins detected in the uterine secretome without phase-specific enrichment. Representative graphs illustrating proteins detected in uterine lavage samples with no phase-specific enrichment across the menstrual cycle. Data are presented as median (IQR). Statistical analysis was performed using the Wilcoxon signed-rank test. NS, not statistically significant. CCL, chemokine ligand; CXCL, chemokine (C-X-C motif) ligand; FGF2, fibroblast growth factor 2; HGF, hepatocyte growth factor; IL, interleukin; PDGF-BB, platelet-derived growth factor BB; SDF-1 alpha, stromal cell-derived factor 1-alpha.

**Supplemental Figure 3.**
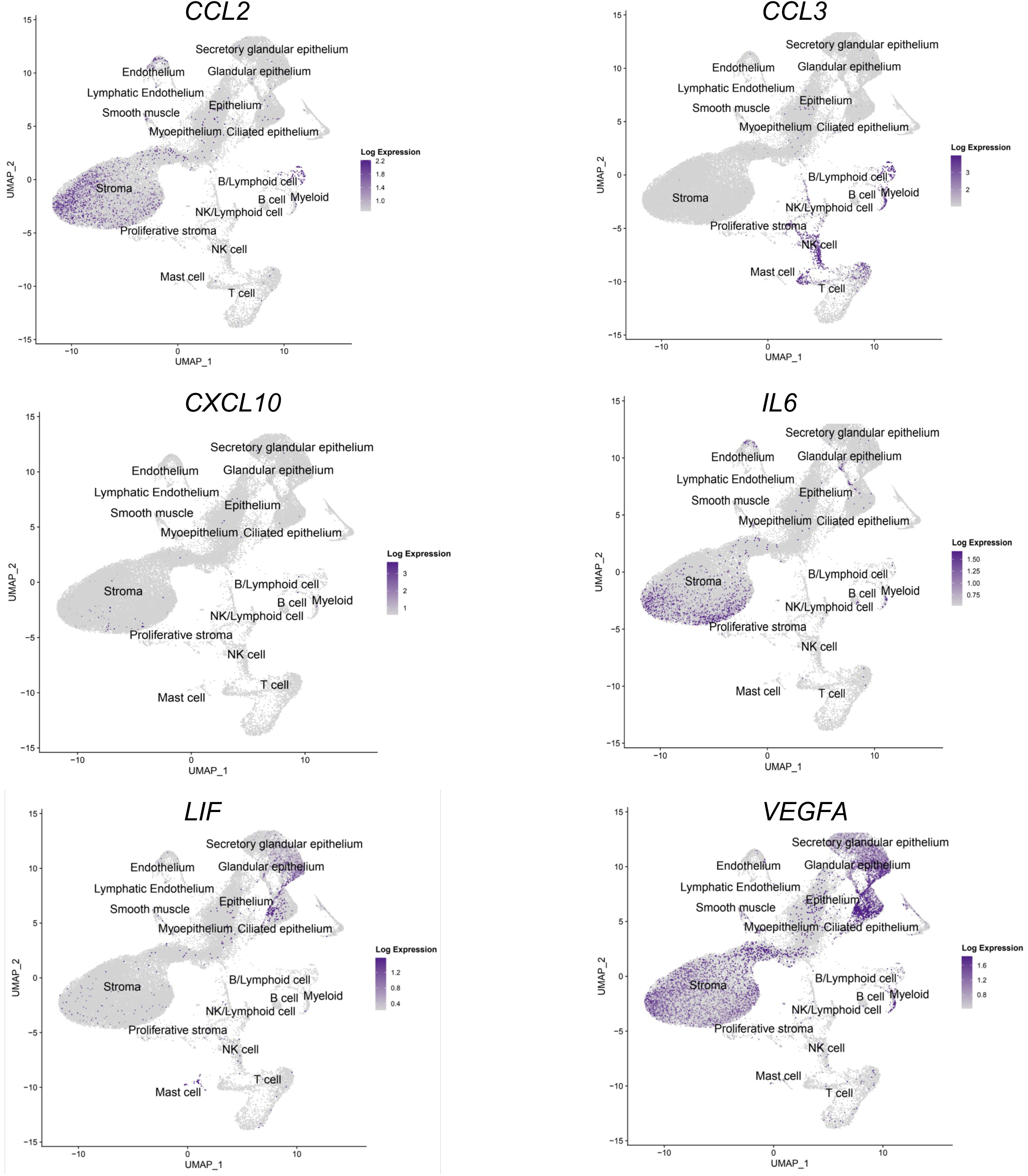
mRNA expression of proteins enriched in the secretome during the window of implantation. UMAP visualization of the six factors expressed and enriched during the mid-secretory phase in both uterine lavage samples and single-cell RNA sequencing. Average score expression is represented by color intensity. CCL, chemokine ligand; CXCL, chemokine (C-X-C motif) ligand; IL, interleukin; LIF, leukemia inhibitory factor; VEGF, vascular endothelial growth factor.

**Supplemental Table 1.** BCA protein concentrations. Total protein concentration in uterine lavage samples measured by BCA assay. BCA, bicinchoninic acid; µg/mL, micrograms per milliliter.

| Supplemental Table 1. BCA Protein Concentrations |  |  |  |
| --- | --- | --- | --- |
| Subject | ID | Cycle Phase | µg/mL |
| 4* | 7 | Proliferative | 203.8 |
|  | 8 | Mid-secretory | 203.8 |
| 5* | 9 | Proliferative | 753.8 |
|  | 10 | Mid-secretory | 953.8 |
| 6 | 11 | Proliferative | 4033.8 |
|  | 13 | Mid-secretory | 1382 |
| 7 | 12 | Proliferative | 1037 |
|  | 21 | Mid-secretory | 453.8 |
| 8 | 14 | Proliferative | 2097.4 |
|  | 25 | Mid-secretory | 397.5 |
| 9 | 16 | Proliferative | 2875 |
|  | 18 | Mid-secretory | 1937.5 |
| 10* | 19 | Proliferative | 1797 |
|  | 22 | Mid-secretory | 1825.5 |
| 11 | 20 | Proliferative | 1010 |
|  | 26 | Mid-secretory | 677.5 |
| 12 | 23 | Proliferative | 2857.5 |
|  | 24 | Mid-secretory | 1062.5 |
| 13 | 27 | Proliferative | 512.5 |
|  | 29 | Mid-secretory | 377.5 |
| 14 | 28 | Proliferative | 370.5 |
|  | 30 | Mid-secretory | 320.5 |
| 15 | 31 | Proliferative | 887.5 |
|  | 32 | Mid-secretory | 330 |

